# Higher levels of narrativity lead to similar patterns of posterior EEG activity across individuals

**DOI:** 10.1101/2022.09.23.509168

**Authors:** Hossein Dini, Aline Simonetti, Enrique Bigne, Luis Emilio Bruni

## Abstract

The focus of cognitive and psychological approaches to narrative has not so much been on the elucidation of important aspects of narrative, but rather on using narratives as tools for the investigation of higher order cognitive processes elicited by narratives (e.g., understanding, empathy, etc.). In this study, we work toward a scalar model of narrativity, which can provide testable criteria for selecting and classifying communication forms in their level of narrativity. We investigated whether being exposed to videos with different levels of narrativity modulates shared neural responses, measured by inter-subject correlation, and engagement levels. Thirty-two participants watched video advertisements with high-level and low-level of narrativity while their neural responses were measured through electroencephalogram. Additionally, participants’ engagement levels were calculated based on the composite of their self-reported attention and immersion scores. Results demonstrated that both calculated inter-subject correlation and engagement scores for high-level video ads were significantly higher than those for low-level, suggesting that narrativity levels modulate inter-subject correlation and engagement. We believe that these findings are a step toward the elucidation of the viewers’ way of processing and understanding a given communication artifact as a function of the narrative qualities expressed by the level of narrativity.

## Introduction

Attention is involved in all cognitive and perceptual processes [1]. To some degree, an attentive state toward an external stimulus implies the silencing of internally oriented mental processing [2]. Sufficiently strong attentive states can hamper conscious awareness of one’s environment and oneself [3]. In contrast to neutral stimuli, emotional stimuli attract greater and more focused attention (see [4] for a review of the topic). As stories can generate and evoke strong feelings [5], the pathos implicit in narratives could be considered attention-seeking stimuli. In fact, people tend to engage emotionally with stories [5]. Attentional focus to a narrative stimulates complex processing [6], with narratives inducing “emotionally laden attention” [2]. In the narrative domain, attentional focus and the sense of being absorbed into the story are part of narrative engagement [3]. Indeed, researchers have suggested that narratives’ inherent persuasiveness is related to the feelings of immersion they evoke, a phenomenon that has been termed “transportation effects” [7]. In this context, “transportation” indicates a combination of attention, feelings, and imagery where there is a convergent process of different perceptual, cognitive and affective systems and capacities to the narrative events [7]. During narrative comprehension, involuntary autobiographical memories triggered by the story in question appear to impair attention only momentarily [8]. This is not the case for either internally or externally generated daydreaming or distraction [8]. Thus, narrative engagement can be hampered when thoughts unrelated to the narrative arise [3]. In addition, the degree to which audiences engage with a narrative varies based on delivery modality (e.g., audio or visual) [9], during narrative comprehension [10], and across individuals [11, 12]. Nonetheless, narratives are naturally engaging [12], and they reflect daily experiences, making them potentially useful devices for understanding cognitive processes such as engagement.

The use of simplified and abstract stimuli has been common in cognitive neuroscience for decades (e.g.: [13]). Although such stimuli enable highly controlled experiments and isolation of the study variables, they tend to lack ecological validity. Hence, recent studies have employed stimuli that simulate real-life situations, including narratives, termed “naturalistic stimuli” (see [12] for a review of the topic). From this perspective, advertising is considered a naturalistic stimulus that is designed to be emotionally persuasive [12]. Video ads provide narrative content in short period of time (e.g., 30 seconds); hence, advertisers focus on delivering a key message through stories with different narrativity levels. In fact, companies use narrative-style ads because stories captivate, entertain, and involve consumers [14,15]. Researchers have found that, at the neural level, narratives (particularly more structured ones) tend to induce similar affective and cognitive states across viewers [2,10]. However, whether consumers perceive stories in similar ways is of great interest, as this might affect whether they ultimately engage with the advertisement as expected. To investigate the underlying cognitive processes generated by narratives, many studies have used traditional methods such as electroencephalogram (EEG) power analysis [16]. Though valid, the metrics provided by traditional methods may not be ideal when considering narrative as a continuous stimulus, given that such methods could generally require stimulus repetition. In addition, they do not capture how the same information is processed across individuals.

Inter-subject correlation (ISC) is an appropriate neural metric for investigating shared neural responses, especially when using naturalistic stimuli. This data-driven method assumes the occurrence of common brain reactions to a narrative, which improves the generalizability of the findings. By correlating neural data across individuals, this metric can identify localized neural activities that react to a narrative in a synchronous fashion (i.e., in a time-locked manner) [17]. ISC is well-suited to analysis of both functional magnetic resonance (fMRI) [18] and EEG [19,20] data. ISC obtained using naturalistic stimuli has been used to investigate episodic encoding and memory [10,21–23], social interaction [24], audience preferences [25], information processing [26], narrative comprehension [27], and, similar to this study, attention and engagement [2,10,11,19,28,29].

ISC calculated from EEG data has been shown to predict levels of attentional engagement with auditory and audio-visual narratives [11,29]. Cohen et al. [29] found that neural engagement with narratives measured through ISC was positively correlated with behavioral measurements of engagement, including real-world engagement. Attention was also shown to modulate (attended) narrative processing at high levels of the cortical hierarchy [26]. Ki et al. [11] found that ISC was weaker when participants had to concurrently perform mental arithmetic and attend to a narrative than when participants only attended to the narrative. This was especially the case when participants attended to audiovisual narratives compared to auditory-only narratives. They also reported that the ISC difference between the two attentional states, as well as its magnitude, was more pronounced when the stimulus was a cohesive narrative than when it was a meaningless one. Moreover, they demonstrated that audiovisual narratives generated stronger ISC than audio-only narratives. In line with this finding, a recent fMRI study indicated that sustained attention network dynamics are correlated with engagement while attending audiovisual narratives [10]. However, this is not the case for audio-only narratives. Furthermore, the authors found higher interpersonal synchronization of default mode network activity during moments of higher self-reported engagement with both audio-visual and audio-only narratives. Finally, ISC was observed to diminish when viewers watched video clips for the second time [2,19,28], which was supported by a study showing lower ISC when viewers were presented with a scrambled narrative for the second time compared to the first time [27].

In summary, while some previous studies have examined the relationship between ISC and narrative engagement (or its correlate, attention) [2,10,11,26,28,29], they assumed that “being a narrative” is an either-or quality. For example, a cohesive narrative is considered a narrative, whereas a meaningless narrative (e.g., a scrambled narrative) is not considered one. From the narratological perspective, there has been increased interest in “narrativity” as a scalar property; that is, artifacts (pictures, videos, stories, and other forms of representation) may have different degrees of narrativity. In other words, narrative may be a matter of more-or-less rather than either-or [30]. Therefore, how different levels of narrativity affect narrative engagement remains an open question. Moreover, whether narrativity levels lead to different degrees of ISC, and whether it is possible to predict narrativity level based on ISC values, has not yet been explored. Therefore, our main research question for this study is as follows: does narrativity level (high [HL] vs. low [LL]) modulate self-reported engagement and ISC? To answer this question, we asked 32 participants to watch 12 narrative, real video ads twice each (with sound removed) varying in their degree of narrativity. We defined “narrativity level” as the degree to which an artifact is perceived to evoke a complete narrative script [30]. Essentially, this refers the degree to which a story seems complete in all its structural elements, such as the presence of defined characters in a stipulated context, clear causal links between events, and a sense of closure resulting from the intertwining of these events [30]. While participants watched the videos, we recorded their EEG brain signals to investigate shared neural responses to the video ads across participants; that is, we investigated the interpersonal reliability of neural responses (expressed by ISC). We further assessed how narrativity level affected self-reported engagement with the ads. Thus, we asked participants to rate how much each ad captured their attention and how immersed in it they felt. Furthermore, we examined how engagement and ISC differ for video ads watched for the first time versus the second time. Based on the aforementioned studies, which explore how narratives engage individuals and indicate that ISC levels are higher during moments of higher narrative engagement and, we postulate that HL video ads will have higher ISC than LL video ads and that self-reported engagement will be higher for HL ads than for LL ads. Moreover, we expect that both engagement and ISC will be higher the first time the videos are seen compared to the second time.

## Results

### Online validation test

Of the 156 participants who completed the online test, we considered 124 responses to be valid as the remaining participants answered too fast or failed to answer the attention question correctly. Results of the online test confirmed that the general public also perceived ads in the HL category as having a high narrativity level (*M_HL_* = 70.95, *SD_HL_* = 10.79) and adds in the LL category as having a low narrativity level (*M_LL_* = 47.71, *SD_LL_* = 13.52; see Fig 5-e in the materials and methods section). All six of the HL ads received higher scores than any of the six LL ads. An independent sample t test confirmed that the difference between the two categories was statistically significant (*t*(61) = 7.554, *p* < .001).

### Narrativity level modulates self-reported engagement

High levels of focused attention and high levels of immersion (feelings of transportation) can indicate high levels of narrative engagement. Thus, we calculated our engagement metric by averaging attention and immersion scores. We first report our descriptive analysis of the attention, immersion, and engagement scores for each of the four conditions (HL–PC, HL–VR, LL–PC, and LL–VR; see “materials and methods” section for abbreviations), then report results of the statistical analysis conducted on engagement scores. Attention scores for the PC conditions were as follows: *M_HL-PC_* was 57.03 (*SD_HL-PC_* = 14.76), and *M_LL-PC_* was 44.16 (*SD_LL-PC_* = 13.92). The attention scores for the VR conditions were as follows: *M_HL-VR_* was 56.26 (*SD_HL-VR_* = 14.19), and *M_LL-VR_* was 46.28 (*SD_LL-VR_* = 14.41). Fig 1-a shows the average attention scores in each of the four conditions. Immersion scores for the PC conditions were as follows: *M_HL-PC_* was 54.26 (*SD_HL-PC_* = 13.28), and *M_LL-PC_* was 42.53 (*SD_LL-PC_* = 15.09). Immersion scores for the VR conditions were as follows: *M_HL-VR_* was 53.67 (*SD_HL-VR_* = 10.81), and *M_LL-VR_* was 44.56 (*SD_LL-VR_* = 14.15). Fig 1-b shows the average immersion scores in each of the four conditions. Engagement scores for the PC conditions were as follows: *M_HL-PC_* was 55.65 (*SD_HL-PC_* = 13.54), and *M_LL-PC_* was 43.35 (*SD_LL-PC_* = 14.04). Engagement scores for the VR conditions were as follows: *M_HL-VR_* was 54.97 (*SD_HL-VR_* = 12.16), and *M_LL-VR_* was 45.42 (*SD_LL-VR_* = 14.00). Results of the 2 × 2 repeated measures ANOVA showed a main effect of narrativity level (*F(1,28)* = 19.779, *p* < .001). Post-hoc analysis showed that participants engaged more with HL ads than with LL ads in both PC (*F(28,1)* = 17.03, *p* < .001) and VR (*F(28,1)* = 20.320, *p* < .001) conditions. See Fig 1-c.

**Fig 1:**
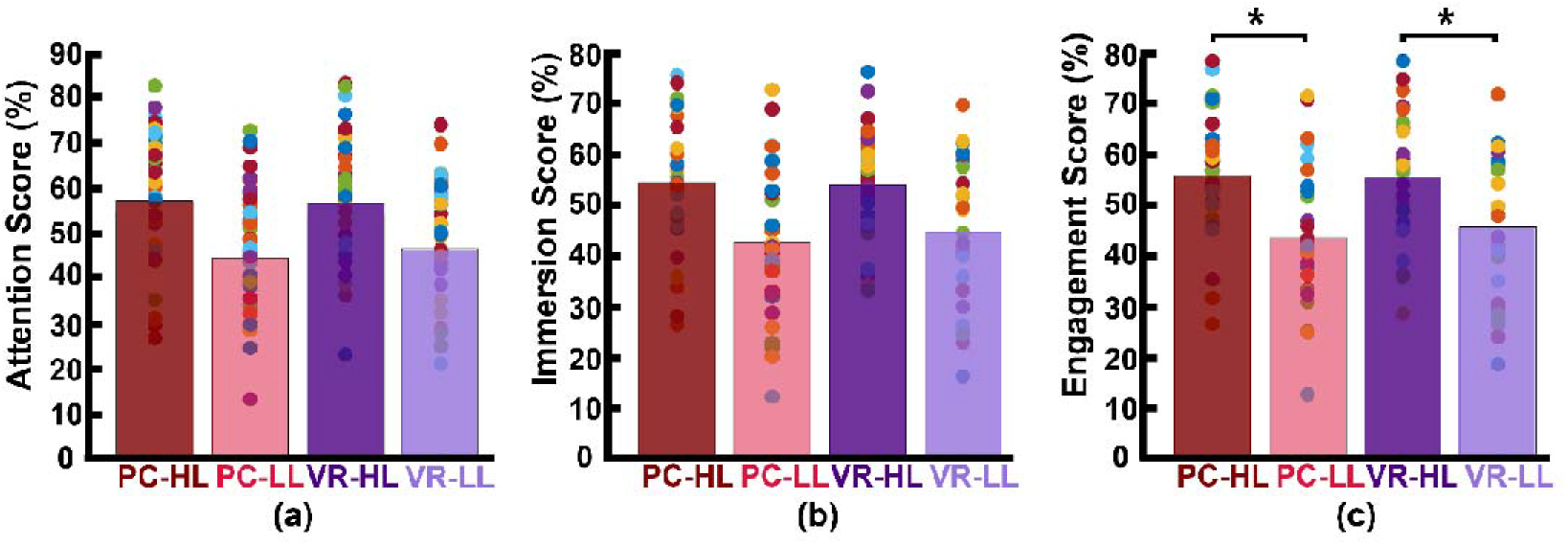
Participants’ self-reported scores. Each bar is the average of the scores across participants and across corresponding video ads. The colored dots represent the score for each participant averaged across the corresponding videos. **(a)** Self-reported attention scores. **(b)** Self-reported immersion scores. **(c)** Engagement scores (average of attention and immersion scores). This shows that the HL engagement scores were significantly higher than the LL engagement scores in both PC (*p* < .001) and VR (*p* < .001) conditions.

### Narrativity level modulates posterior inter-subject correlation

We applied the same statistical analysis used for the self-reported data to assess whether our neurological metric (ISC) changed according to narrativity level. To do so, we used summed ISC over the first two strongest components (see Materials and Methods section) as a representative of neural reliability. Results of the 2 × 2 repeated measures ANOVA showed a main effect of narrativity level, where ISC was higher when participants watched the HL ads than when they watched the LL ads (*F (1,28)* = 4.467, *p* = .044). Although the interaction effect was not significant, we then conducted post-hoc analysis to determine whether this difference occurred in the PC or VR condition. Results of the post-hoc analysis showed that ISC was significantly higher for the HL ads (*M_HL_* = 5.47 × 10^−3^, *SD_HL_* = 2.97 × 10^−3^) than for the LL ads (*M_LL_* = 3.48 × 10^−3^, *SD_LL_* = 2.51 × 10^−3^) in the PC condition (*F (28,1)* = 5.266, *p* = .017). In the VR condition, the ISC of HL ads (*M_HL_* = 8.72 × 10^−3^, *SD_HL_* = 4.62 × 10^−3^) was still higher than that of LL ads (*M_LL_* = 8.35 × 10^−3^, *SD_LL_* = 3.92 × 10^−3^), but this difference was not significant (*F (1,28)* = 0.226, *p* = .638). See Fig 1-a.

Furthermore, to investigate which factors affect neural reliability, we compared the ISC of HL video ads to that of LL video ads separately for each component. We corrected the p-values because we repeated our calculations twice for the two components, and results can be seen in the top part of Fig 2-b. Results showed that, for the first component, the main effect of narrativity is not significant (*F (1,28)* = 1.145, *corrected p* =.294). However, for the second component, the main effect of narrativity was significantly higher in HL video ads than in LL video ads (*F (1,28)* = 5.412, *corrected p* = .045). Post-hoc analysis of the second component revealed that the ISC of HL ads (*M_HL_* = 2.65 × 10^−3^*, SD_HL_* = 3.34 × 10^−4^) was significantly higher than that of LL ads (*M_LL_* = 9.61 × 10^−4^, *SD_LL_* =2.99 × 10^−4^) in the PC condition (*F (1,28)* = 13.676, *corrected p* = .002); however, there was no significant difference between HL and LL ads in the VR condition (*F (1,28)* = 1.056, *corrected p* = .313). In addition, different correlated components had different patterns of scalp activities (Fig 2-b, bottom part). For the first component, activity was distributed throughout the anterior and posterior regions, while in the second component, activity was concentrated in the posterior region. Overall, the second component drove the narrativity level modulation in response to the HL narrative video ads.

**Fig 2:**
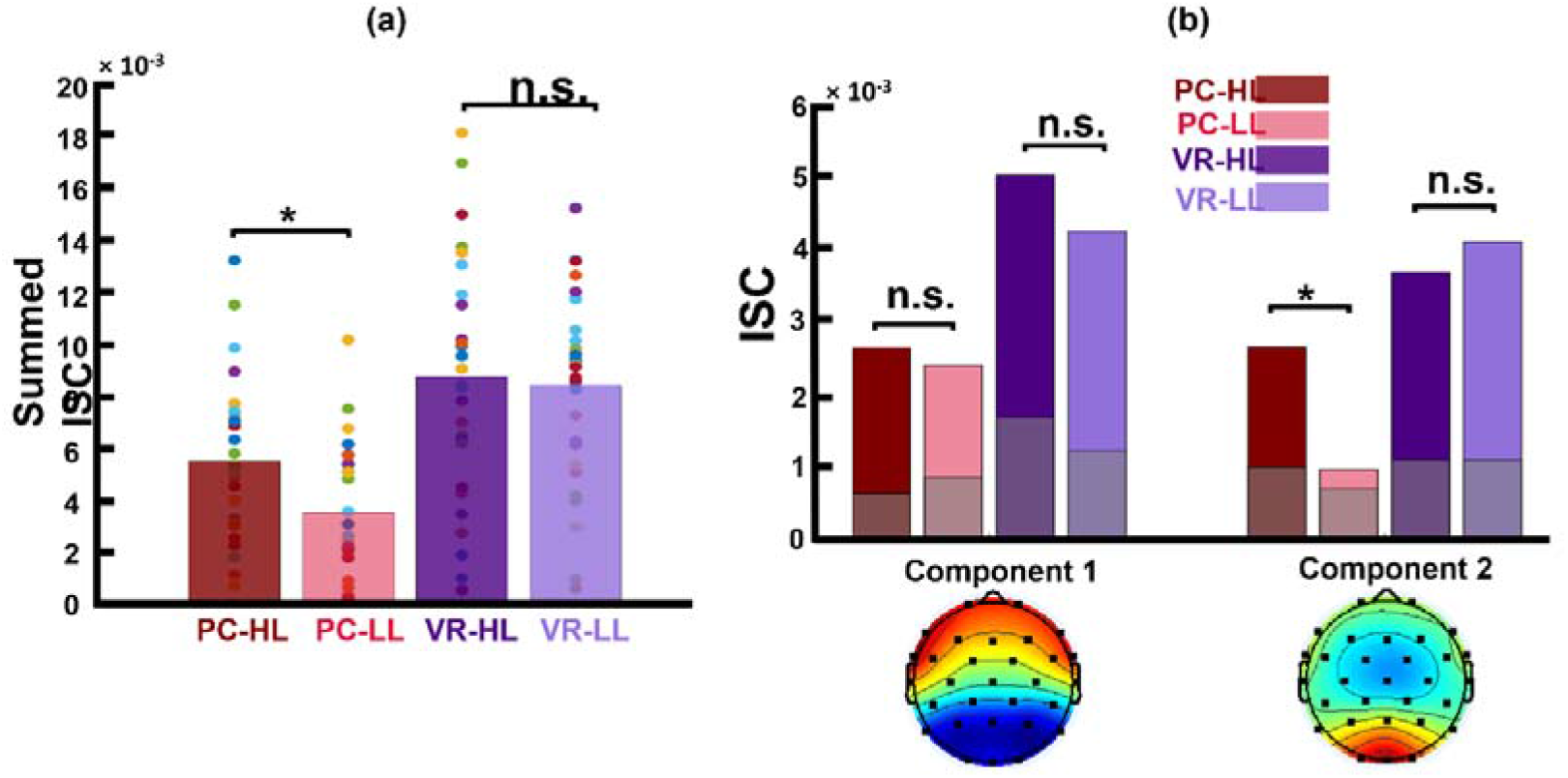
**(a)** Summed ISC over the first two strongest components in four conditions. This shows that, in the PC condition, the ISC of HL video ads was significantly higher than that of LL video ads (*p* = .029), and in the VR condition, the ISC of HL video ads was still higher than that of LL video ads, but not significantly so (*p* = .638). **(b-top)** Calculated ISC separately for each component and within each condition. This shows that, for the first component, the difference between HL and LL video ads is not significant in either the PC or the VR condition. However, in the second component, the ISC of HL video ads was significantly higher than that of LL video ads (*p* = .002) in PC condition, but not in the VR condition (*p* = .313). The gray bars show the average calculated chance level for each condition **(b-bottom)** Scalp activity of first two strongest components. Activity in component 1 is throughout the anterior and posterior regions, while in component 2, activity is concentrated in the posterior region. In both figures, the colored dots represent the calculated ISC values for each participant.

Previous studies found a direct link between engagement and ISC. Thus, to investigate whether the narrativity level modulated ISC through engagement, we conducted a mediation analysis: narrativity level → engagement → ISC of the second component using the combined data in PC and VR. The results confirmed the total effect of narrativity level on ISC (total effect = 0.64 × 10^−3^, *t(26)* = 2.305, *p* = .029, 95% C.I. [0.07 × 10^−3^ × 1.21 10^−3^]). In addition, they showed a direct effect of narrativity level on ISC scores: that is, an effect when controlling for engagement levels (direct effect = 0.80 × 10^−3^, *t(26)* = 2.198, *p* = .037, 95% C.I. [0.05 × 10^−3^ × 1.56 10^−3^]), and no indirect effect of narrativity level on ISC scores through engagement (indirect effect = −0.16 × 10^−3^, 95% C.I. [-0.63 × 10^−3^, 0.14 × 10^−3^]).

### Viewing order modulates self-reported engagement but not inter-subject correlation

To evaluate whether viewing order affects engagement scores and ISC, we implemented the abovementioned statistical procedure with narrativity level and viewing order as independent variables. Participants watched each video twice: once in one medium and once in the other medium. The medium that was used first was counterbalanced across participants. For the analysis, we created a group for the first viewing and a group for the second viewing, regardless of the medium used. In both of these groups, participants watched the same videos, including both HL and LL video ads. We first report the resulting engagement scores and ISC scores.

Regarding engagement scores, the average and standard deviation for the first viewing group were as follows: *M_HL_* was 57.71 (*SD_HL_* = 2.10), and was 45.42 (*SD_LL_* =2.65). For the second viewing group, these figures were as follows: *M_HL_* was 52.90 (*SD_HL_* = 2.57), and *M_LL_* was 43.36 (*SD_LL_* =2.56; see Fig 3-a). Results of this statistical analysis revealed a significant main effect of viewing order. Further post-hoc analysis indicated that this main effect was driven by the HL category, for which videos viewed first received significantly higher engagement scores than those viewed second (*F(1,28)* = 12.784, *p* = .001). However, in the LL condition, there was no significant difference between the first and second viewing groups (*F(1,28)* = 1.686, *p* = .205). Moreover, there was a main effect of narrativity level: of videos in the first viewing group, HL videos received significantly higher engagement scores than LL videos (*F(1,28)* = 23.047, *p* < .001); this was also the case for videos in the second viewing group (*F(1,28)* = 13.391, *p* =.001).

**Fig 3:**
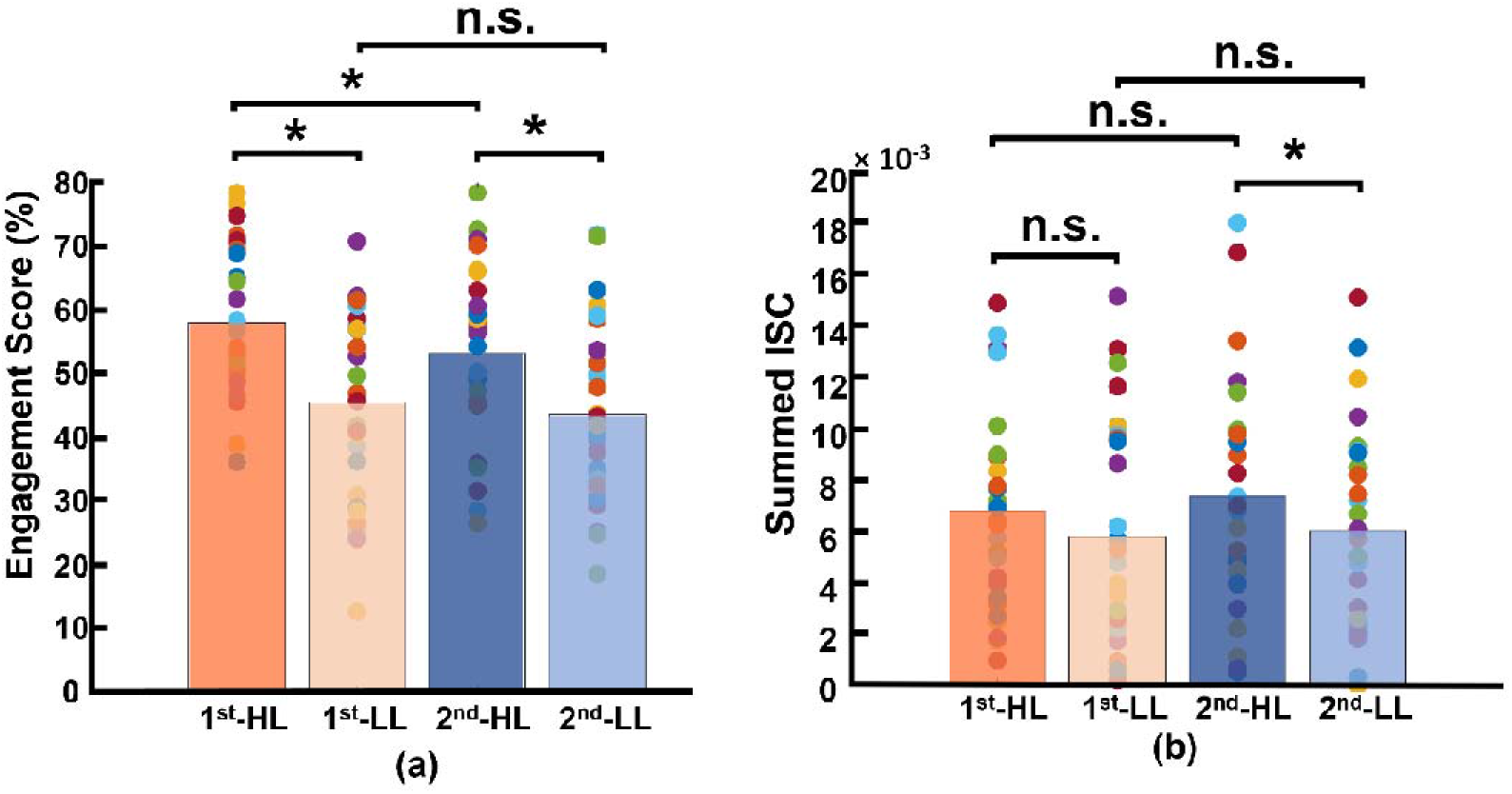
Results of viewing order (first and second) considering narrativity level (HL and LL). **(a)** Results of engagement scores, where for videos in the HL condition, engagement scores upon first viewing were significantly higher than those upon second viewing. Moreover, HL videos received significantly higher engagement scores than LL videos upon both first and second viewings. (**b)** Results for calculated ISC of viewing order, where the only significant difference was between the second viewing of HL videos and that of LL videos.

Regarding ISC, the average and standard deviation for videos in the first viewing group were as follows: *M_HL_* was 6.92 × 10^−3^ (*SD_HL_* = 0.74 × 10^−3^), and *M_LL_* was 5.97 × 10^−3^ (*SD_LL_* = 0.81 ×10^−3^). For the second viewing group, these figures were as follows: *M_HL_* was 7.27 × 10^−3^ (*SD_HL_* = 0.82 × 10^−3^), and *M_LL_* was 5.88 × 10^−3^ (*SD_LL_* = 0.71 × 10^−3^; see Fig 3-b). Results of this statistical analysis showed no significant main effect of viewing order. Further post-hoc analysis showed that the HL video ads received higher ISC scores than LL video ads upon second viewing (*F(1,28)* = 4.411, *p* = .045), but not upon first viewing (*F(1,28)* = 1.144, *p* = .294).

### ISC-HL

We calculated ISC-HL to test if it was possible to determine whether a single subject was watching an HL video ad or an LL video ad based on their neural activity. We calculated the projections based only on data from the HL group. These projections represent the overall brain activity of all participants attending to HL videos. Based on these projections, we calculated the ISC-HL for participants from each group (HL and LL) separately. We hypothesized that the brain activity of participants from the HL group would be more similar to the overall activity of all participants in the HL group (meaning higher ISC-HL) and that the brain activity of participants from the LL group would be less similar to the overall activity of all participants in the HL group (meaning lower ISC-HL). Fig 4-a and Fig 4-b display the calculated ISC-HL in both VR and PC conditions. As expected, the neural responses of 23 (76.6%) participants were much more similar to those of the HL group (had higher ISC-HL) when they attended to HL videos than when they attended to LL videos in the PC condition (Fig 4-a). In the VR condition, the neural responses of 15 (50%) participants showed higher ISC-HL while attending to HL videos than while attending to LL videos (Fig 4-b). To assess the classification performance of the support vector machine classifier with 10-fold crossvalidation for the two groups (attended to HL videos vs. attended to LL videos), we calculated the area under the curve (AUC). To do so, we first calculated the actual AUC value by calculating the projections using actual labels, then randomly shuffled the labels 1,000 times. For each iteration, we calculated new projections and computed the AUC. The average AUC over 1,000 iterations was the chance-level AUC. Given ISC-HL as the predictor of narrativity level, the classifier shows above-chance performance in the PC condition but not in the VR condition (Figs 4-c and 4-d). In the PC condition, the actual AUC was .637, while the chance-level AUC was .548. In the VR condition, the actual AUC was .507, while the chance-level AUC was .543. Therefore, in the PC condition, the ISC-HL was able to predict the narrativity level based on neural responses (measured via EEG) to the stimuli. This means that the model significantly predicted whether the participants were attending to HL or LL narrativity. The level of significance was not as strong as the previously reported accuracy for predicting attentional states [11]; however, this could be due to explicit differences between experimental conditions (participants were made to count backward during the task to diminish their attentional state). This finding is the first step toward predicting exposure to different levels of narrativity based solely on the ISC of neural activity evoked by stimuli, which leads to a better comprehension of the brain mechanism that processes narrativity levels. Further studies with different designs should be conducted with a more specific focus on predicting narrativity level.

**Fig 4:**
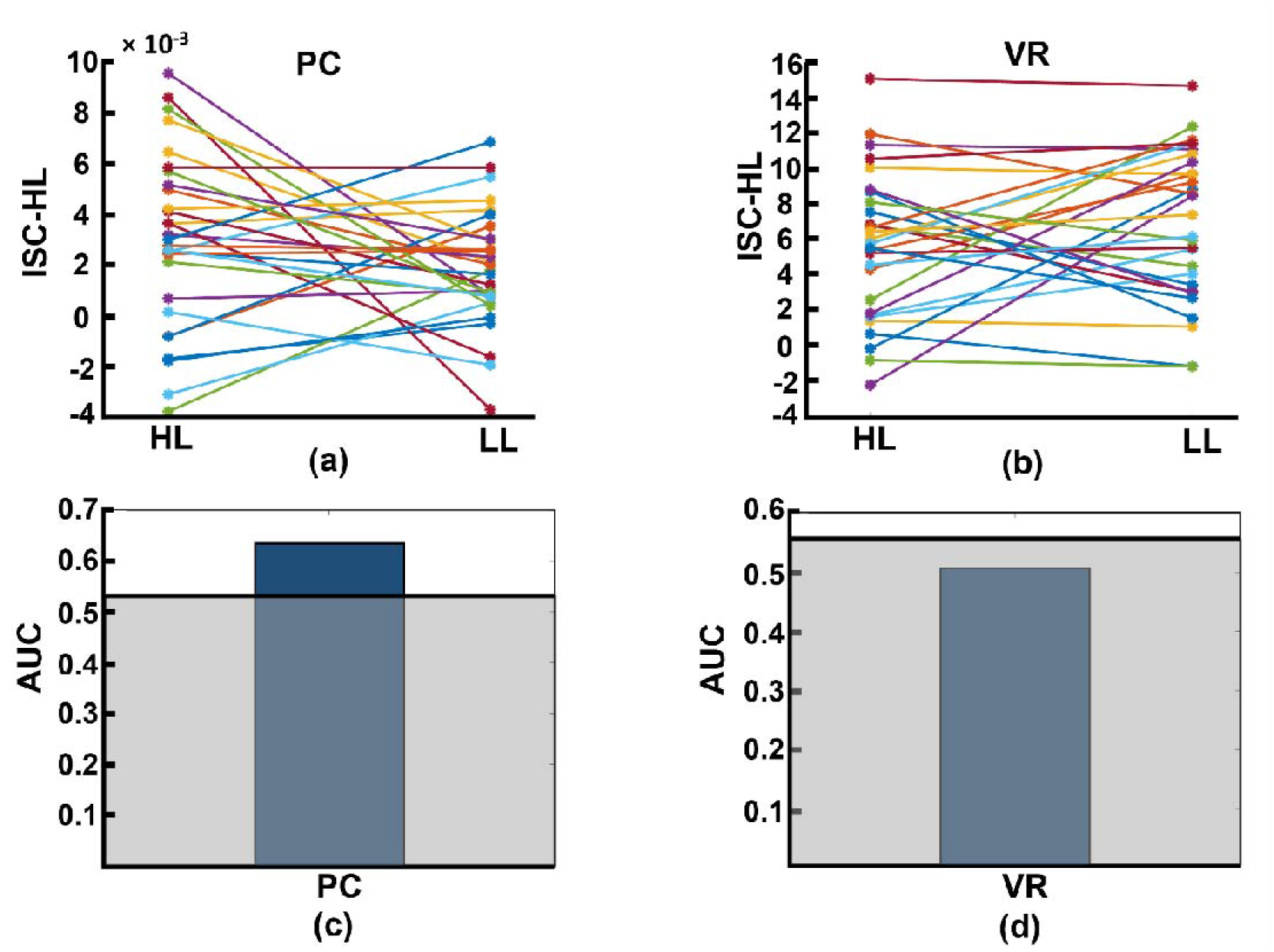
**(a) and (b)** Results for calculated ISC-HL for each participant in the PC and VR conditions. Lines and markers of each color show the ISC-HL of one participant in the HL and LL conditions. **(a)** This shows that 76.6% of participants had higher ISC-HL when they attended to HL video ads than when they attended to LL video ads. This shows that most of the participants who were exposed to HL video ads showed higher similarity to the trained model (which is derived from data of HL condition) than those who are exposed to LL video ads. **(b)** This relates to the same evaluation as (a), but in the VR condition, where 50% of participants had higher ISC-HL. **(c)** and **(d)** AUC of classification performance for predicting exposure to HL and LL video ads in PC and VR conditions, respectively. The gray area shows the chance-level performance. **(c)** shows that the AUC of classification based on participants’ neural activity in the PC condition is higher than chance level, while **(d)** shows that the AUC of classification based on participants’ neural activity in VR condition is below chance level.

**Fig 5:**
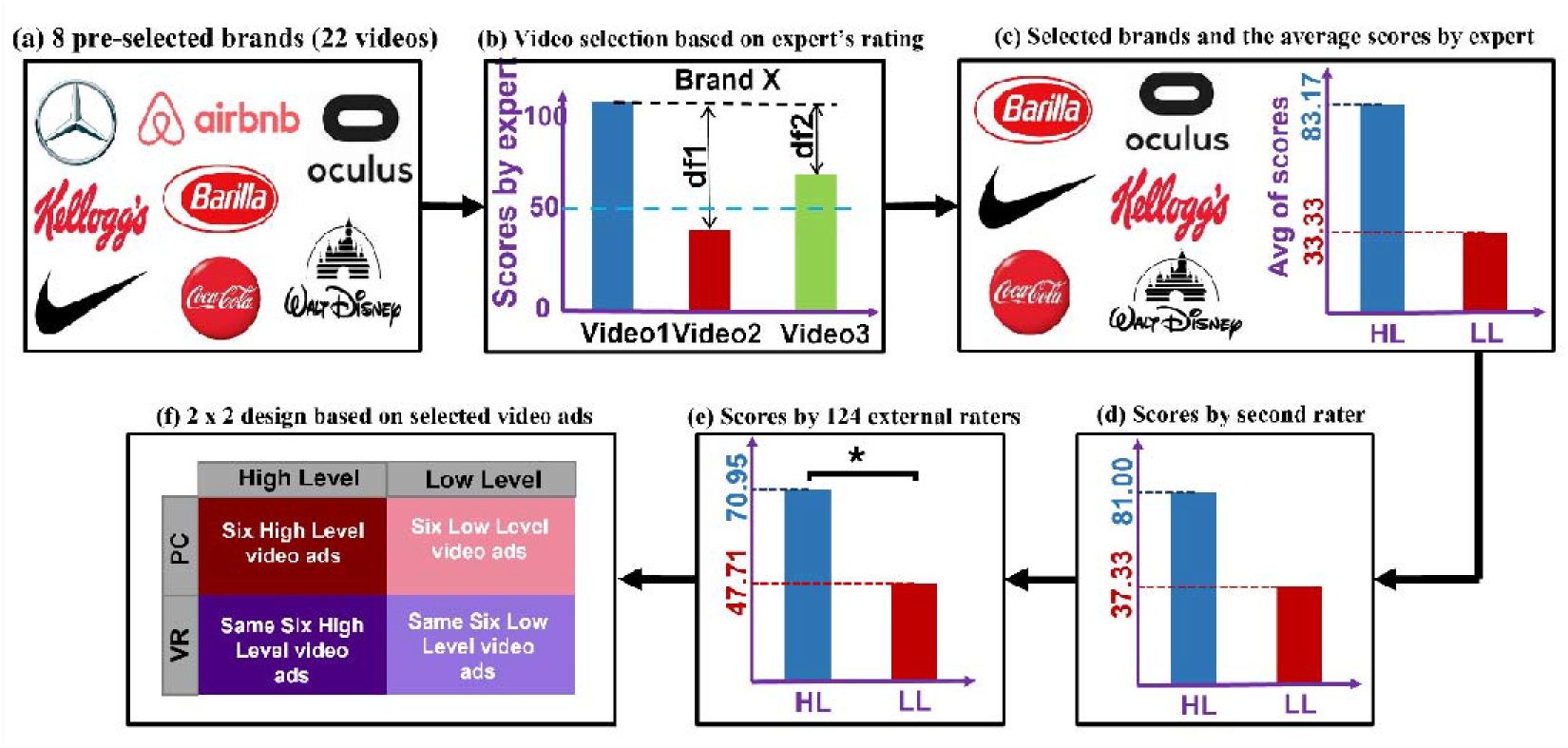
Video ads selection procedure, division into HL and LL categories, results of the ratings, and the resulting design. **(a)** The eight preselected brands from which we chose 22 video ads. **(b)** Process of selecting videos based on expert’s rating. Suppose that, from brand X, we preselected three videos that were rated by the expert. The videos that had the maximum difference (df1) in score, for which one score was > 50 and the other was < 50, were selected as the final videos from brand X. **(c)** The six selected brands with average scores. For each brand, the video with a higher score was assigned to the HL category (six videos) and the video with a lower score was assigned to the LL category (six videos). **(d)** Average of the second expert’s ratings across the six selected videos. **(e)** Average scores (across participants and stimuli) for narrativity level, rated by 124 participants. An independent sample t test showed that the narrativity scores of the HL videos were significantly higher than those of the LL videos (*p* < .001). **(f)** The 2 × 2 experiment design consisted of the following conditions: HL–PC, HL–VR, LL–PC, and LL–VR.

## Discussion

In this study, we tested whether different levels of narrativity (HL vs. LL) lead to differences in information processing reliability (represented by ISC), specifically while watching video ads. Furthermore, we evaluated whether different levels of narrativity cause differences in self-reported engagement ratings. To this aim, we presented HL and LL video ads to 32 participants while collecting their EEG signals. In addition, for each video ad, participants self-reported their levels of attention and immersion, which we considered two core factors of engagement. We calculated the ISC of each participant by calculating the similarity of their correlated components to those of the participant pool. One advantage of ISC analysis is that it does not require stimulus repetition; this is advantageous because such repetition causes decreased attention and engagement [2,11]. As expected, our results showed that both calculated ISC and engagement scores of HL video ads were significantly higher than those for LL video ads, suggesting that narrativity level modulates ISC and engagement. However, the modulation of ISC was not mediated by the degree of engagement with the narrative.

Previous studies have evaluated the relationship between narrative engagement—or its core component, attention—and ISC. Cohen et al. measured self-reported engagement scores and ISC while participants were exposed to naturalistic videos [29]. They found that more engaging videos were processed uniformly in participants’ brains, leading to higher ISC. Consistent with their results, Poulsen et al. reported that a lack of engagement manifests an unreliable neural response (meaning lower ISC), and they introduced EEG-ISC as a marker of engagement [28]. Song et al. investigated whether engagement ratings modulate ISC by evaluating the relationship between continuous self-reported engagement ratings and ISC using continuous naturalistic stimuli [10]. They reported higher ISC during highly narrative engaging moments and concluded that ISC reflects engagement levels. In another study, Dmochowski et al. used short video clips to evaluate attention and emotion using ISC [2]. They found a close correspondence between expected engagement and neural correlation, suggesting that extracting maximally correlated components (ISC) reflects cortical processing of attention or emotion. Finally, Ki et al. investigated whether attentional states modulate ISC for audio and audiovisual narratives, and they concluded that higher attention leads to higher neural reliability across subjects [11]. Inspired by previous studies [2,3,31], we measured narrative engagement by averaging two of its important components, attention and immersion.

Our results indicated that, using ISC (in the PC condition), we were able to significantly discriminate levels of narrativity: the ISC levels of HL video ads were significantly higher than those of LL video ads (Fig 2-a). Consistent with our narrativity level discrimination, self-reported engagement ratings were significantly higher for HL video ads than for LL ones (Fig 1-c). However, in contrast with other studies, we did not find a relationship between engagement levels and ISC. A plausible explanation for this finding is the type of stimulus employed.

Several studies compared ISC scores of interrupted or scrambled narratives with those of non-scrambled narratives. Dmochowski et al. reported higher ISC for non-scrambled narratives than for disrupted or scrambled narratives [2]. In addition, Ki et al. and Poulsen et al. found higher ISC scores for a cohesive narrative than for a meaningless, scrambled narrative [11,28]. These findings demonstrate that scrambled narratives are linked to lower ISC scores. Song et al. used scrambled video clips to identify the moments in which narrative comprehension occurs [27], and they reported that story comprehension occurs when events are causally related to each other. Moreover, they showed that, in such moments, the underlying brain states were mostly correlated across subjects. The stimuli manipulations applied by previous studies (e.g., scrambling the narrative or presenting narratives that lacked causality) created greater differences between experimental conditions than those in our study. While we shared with these studies their interest in the subjects’ engagement and other factors related to the reception of the narrative, we were also interested in investigating the possibility of classifying and discriminating structural characteristics of narrative artifacts. Therefore, we defined the narrativity levels (HL and LL) based on narrative structural elements and properties such as the presence of defined characters in an identifiable context, clear causal links between events, and closure resulting from the intertwining of these events. Our conceptualization of narrativity levels was inspired by Ryan [30].

From this perspective, our results align with those of previous studies, indicating that information processing is highly consistent across subjects when participants are exposed to HL narrativity. In other words, the fact that the ISC of HL video ads is significantly higher than that of LL video ads in the PC condition (Fig 2-a) is a promising indication that EEG-ISC could potentially be used to explore levels of narrativity (and correlative engagement) in different kinds of media artifacts. However, unlike previous studies using stimuli that were “either-or” regarding narrativity possession, our ISC was not mediated by engagement levels. This finding suggests that ISC can represent or capture other cognitive processes beyond engagement. The evaluation of the activity of the first two components separately (Fig 2-b) provides some insights in this regard. Results showed that the second component (in the PC condition) was most reliably modulated by the narrativity level. The second component had the strongest shared activity concentrated in the posterior part of the brain. While previous studies relating ISC to engagement found more widespread ISC (e.g., [2,10,25,29]), strong posterior ISC was linked to shared psychological perspective [32]. Lahnakoski et al. asked participants to take one perspective or another to interpret the events of a movie. When participants watched the movie and adopted the same perspective, posterior ISC was stronger than when they adopted different perspectives. The authors posited that ISC represented a shared understanding of the environment [32]. In our case, high levels of narrativity better immersed participants in the story world compared to low levels. This might have eased participants to take the perspective of the character(s) in HL videos. In addition, videos with low narrativity levels might not have been so successful in leading to similar perspective taking because the story was more fragmented than in HL video ads. However, further investigation is necessary to elucidate why narrativity level didn’t appear to affect ISC in the same way in the VR condition. Chang et al. reported that ISC might change due to fatigue effects for participants in an fMRI scanner [33]. Thus, one possible explanation for the non-significant effect of narrativity level on ISC in the VR condition is fatigue. Wearing the VR headset and the EEG cap for almost 25 minutes might have caused fatigue and discomfort (e.g., related to posture, weight, or itching), and such an effect might have masked the effects of narrativity level on ISC. Another plausible explanation is also related to the VR feature. As VR is an increasingly popular immersive technology, participants’ expectations about the VR modality may have affected their level of attention. Participants may have been disappointed to find a 2D stimulus that perhaps failed to meet their expectations and therefore diminished their active attention.

In the narratology field, there is a substantial, ongoing debate about whether narrative should be considered an either-or property or a scalar property (i.e., a matter of more or less) [30,34]. Therefore, in the last two decades, some narratologists have introduced the notion that different artifacts may have different levels of narrativity [30,35]. It is widely accepted that, even though events in a story do not have to be chronologically ordered, the sequence of the events must nonetheless follow a narrative logic if closure is to be achieved (this is something that is commonly encapsulated in the distinction between “story” and “discourse”). Therefore, a “scrambled” narrative ceases to be a narrative; instead, it is a series of unrelated events, lacking an internal temporal logic. If there is no discernable, recoverable chronological order of connected events, the sequence could hardly be considered a narrative. Although the previous studies presented in our literature review provided the methodological bases for our study, they were not centered on narrative qualities. In our study, we focus on investigating the plausibility of testing a particular expressive artifact for levels of narrativity. Therefore, our stimuli presented two different degrees of narrativity (LL and HL). We showed that small differences in narrativity levels (i.e., between HL and LL) have effects on ISC and engagement similar to the effects of more evident differences between such levels (e.g., scrambled vs. non-scrambled). However, the underlying reason for differences in ISC between narratives with some degree of narrativity level did not reflect differences in perceived engagement; rather, it seemed related to perspective taking. These findings are a step toward narrative comprehension, especially when considering narrative as a scalar property [36].

To test whether viewing order modulates engagement and ISC, we separated the data into two groups—first and second viewing—and conducted statistical analyses considering narrativity level and viewing order to be independent variables. Domchowski et al. found that attentional engagement decreases when participants watch a stimulus for the second time compared to the first time [2]. Moreover, they reported significantly lower ISC for the second viewing. Ki et al. confirmed and extended their results by declaring that neural responses become less reliable upon second viewing of a stimulus, showing significantly lower ISC [11]. In an fMRI study, Song et al. investigated the effect of viewing order on neural activity, watching scrambled videos twice [27]. Their results replicated the findings of the aforementioned study and other studies [19,28] by showing that neural states across participants were less synchronized when watching the videos, including the scrambled videos, for the second time. However, Chang et al. conducted a combined EEG-magnetoencephalography (MEG) study and reported increased ISC during the second viewing of the stimuli [33]. They declared that participants’ prediction of the story structure increased during the second viewing, resulting in a more similar EEG-MEG activity across participants, and that this contradiction with previous studies might be due to the different time window selected for ISC calculation. Our results showed that engagement level dropped during the second viewing, supporting previous studies [2]. For the HL videos ads, engagement scores for the first viewing were significantly higher than those for the second viewing, although this was not the case for the LL ads (Fig 3-a). The decrease in engagement scores for HL ads but not LL ads might be due to differences intrinsic to the classification of our stimuli. While videos in the HL category included all or almost all narrativity elements [30], those in the LL category included only a few. Thus, we could say that the HL video ads were more storytelling-based than the LL video ads. Therefore, it could be that factors such as suspense, expectation, and suspension of disbelief, which naturally occur when watching stories, are attenuated when the story is viewed for the second time, lowering engagement levels in the case of HL video ads. These same factors are not present or are present to a lesser extent when watching LL video ads, and therefore, engagement levels were not harmed. However, there was no significant difference in ISC between the first and second viewing in either HL or LL video ads, suggesting that viewing order does not modulate ISC. Though the current dataset cannot provide definite answers, these differences in findings might be explained by our study design. First, our participants were exposed to 12 short video ads, a greater number of stimuli than previous studies employed when investigating viewing order [2,11,27,28,33]. Watching 12 videos in sequence might have reduced participants’ memories of details of the stories. Hence, when watching the videos for the second time (although they were perceived as less engaging) there remained a substantial amount of information to be processed, which could have been reflected in the ISC of the second viewing. Another possible factor that might have hindered participants’ short-term memories of the videos and affected ISC levels is the time span and the tasks performed between the two viewings. In our study, the second exposure to the video ads was separated from the first by about 20 min. During those 20 min, participants performed two different tasks for a separate study. Therefore, the current dataset was not able to capture the neural underpinnings of viewing order. Considering the limitations of this study, future studies could be conducted to capture this effect.

## Materials and methods

This study was approved by the local ethics committee (Technical Faculty of IT and Design, Aalborg University) and performed in accordance with the Danish Code of Conduct for Research and the European Code of Conduct for Research Integrity. All participants signed an informed consent form at the beginning of the session, were debriefed at the end of the experiment, and were given a symbolic payment as a token of gratitude for their time and effort. Data collection took place throughout November 2020. This study was part of a larger study; here, we report only the information relevant to the present study.

### Participants and stimuli

We recruited 32 (13 women) right-handed participants of 16 nationalities between the ages of 20 and 37 (*M* = 26.84, *SD* = 4.33). Regarding occupation, 69% were students, 16% were workers, and 15% were both. Regarding educational level, 12% had completed or were completing a bachelor’s degree, and 88% had completed or were completing a master’s degree. We requested that participants not drink caffeine products at least two hours prior the experimental session.

To select the stimuli, we initially pre-selected 22 video ads (eight brands; Fig 5-a) varying in the number of elements that constitute different narrative levels according to Ryan [30]. The narrativity levels of these video ads were then preliminarily rated from 0 to 100 by an expert in the narrative field. Based on these scores, we selected six brands that had the largest difference in score between two of their ads (to maximize the difference in narrativity level between two ads of the same brand), coinciding with an absolute score below 50 for one ad and above 50 for the other ad (Fig 5-b). We then grouped the video ads into two categories: HL and LL narrativity. Although not categorizing the videos—thus considering continuity in narrativity level—would better align with a scalar model of narrativity, our results would suffer from statistical invalidity. Because the videos were real and therefore not created specifically for the study, they had idiosyncratic features separate from the narrativity level (e.g., different scenario, characters, plot). These features could either cover a potential effect of narrativity level or lead to a misattribution of an effect. By categorizing the videos into two narrativity levels, we sought to mitigate the potential effects of those individual features and make the differences in narrativity level more salient. The final selection consisted of six video ads with HL narrativity (*M* = 83.17, *SD* = 10.70) and another six with LL narrativity (*M* = 33.33, *SD* = 10,70), with each of the six brands contributing one video to each of the two groups (Fig 5-c). To validate the categorization of the ads into these two categories, another independent expert in the narrative field rated the narrativity level of the 12 video ads from 0 to 100, and their ratings were consistent with those of first expert (HL: *M* = 81.00, *SD* = 13.70; LL: *M* = 37.33, *SD* = 15.37; Fig 5-d). To assess whether the general public would also perceive a difference in narrativity level between the two categories of videos, we conducted an online validation test using the Clickworker platform (https://www.clickworker.com) with an independent panel of participants, which confirmed that our classification was valid (see the results section). For the online test, a total of 156 participants were assigned to one of four groups. Each group was assigned to watch three video ads, each of which was from a distinct brand. To capture the degree of perceived narrativity in each video ad, participants were asked to answer five questions used by Kim et al. [37], such as “the commercial tells a story” and “the commercial shows the main actors or characters in a story.” All questions were rated from 0 (strongly disagree) to 100 (strongly agree). Average responses to these five questions showed whether the videos ads were indeed perceived to be in the HL (higher scores) or LL (lower scores) category. The responses confirmed that our initial categorization of the videos was valid (see results and Fig 5-e).

The stimuli therefore comprised 12 2D video ads from six brands (Barilla, Coke, Disney, Kellogg’s, Nike, and Oculus), with two video ads from each brand, one for each narrativity level. We selected different brands to mitigate the influence of brand and product category. The ads were real commercials retrieved from YouTube. We removed the audio and edited some of the videos slightly to adjust the length (which varied from 57 to 63 seconds). We selected non-verbal narratives in video format because motion picture narratives are less susceptible to interindividual differences and generate more homogeneous experiences across individuals than verbal (oral) narratives [38]. This may be because the visual images are directly related to the narrative content, which reduces personal interpretations of the story [9].

### Design, data collection, and task

We conducted a 2 × 2 full factorial within-subjects study with two levels of narrativity (high level [HL] vs. low level [LL]) and two media (computer screen [PC] vs. virtual reality [VR]). The study included four conditions: six HL video ads presented on a PC (HL–PC), six HL video ads presented in VR (HL–VR), six LL video ads presented on a PC (LL–PC), and six LL video ads presented in VR (LL–VR). See Fig 5-f.

Initially, the EEG device (32-channel, 10-20 system) was placed on each participant’s scalp, and the impedance of the active electrodes was set to less than 25 kΩ using a conductive gel, according to the manufacturer’s guidelines. The signals were recorded at a sampling rate of 500 Hz using Brain Products software. The HTC Vive Pro VR headset was placed on top of the EEG electrodes, and the impedance of the electrodes was checked again. The task comprised two sessions of approximately 25 min each, separated by a 20 min interval. During one session, participants watched ads on a PC; during the other session, they watched ads in VR. During each session, participants watched all of the video ads and answered a questionnaire after each. A two-second fixation cross was shown before each new video. The videos were displayed first on a PC for half of the participants. The video presentation order was counterbalanced across participants but the same across media. The ten-item questionnaire included two questions of interest for this study: (i) “this commercial really held my attention” and (ii) “this ad draws me in”; both were scored from 0 (strongly disagree) to 100 (strongly agree). We retrieved these questions from the “being hooked” scale[39], and they were meant to capture consumers’ sustained attention to the advertisement [39]. Our engagement metric was calculated based on the combined average scores of these two questions.

### Preprocessing

We performed all preprocessing and processing steps using Matlab R2020b (The Math Works, Inc) with in-house codes and tools from the FieldTrip 20210128 (http://fieldtriptoolbox.org) and EEGLAB 2021.0 (https://eeglab.org/) toolboxes. To remove high and low frequency noise, we applied a third-order Butterworth filter with 1–40 Hz cut-off frequencies to the raw data. Next, we detected bad channels using an automated rejection process with voltage threshold ±500 μV and, after confirmation by an expert, rejected them from the channel list. We then interpolated the removed channels using the spherical spline method based on the activity of six surrounding channels in the FieldTrip toolbox. The average number of rejected channels per participant was 1.92 ± 1.48. One participant was excluded from the analyses due to having more than five bad channels, and two participants were excluded due to missing trials. Subsequently, we segmented the filtered EEG data corresponding to the 12 video ads and concatenated the data. We then excluded the EEG data corresponding to the time when participants were answering the questionnaires. Next, we conducted independent component analysis on the concatenated data to remove remaining noise. We estimated source activity using the second-order blind identification method. We then identified eye-related artifacts and other noisy components, which were confirmed by an expert and removed from the component list. Applying the inverse independent component analysis coefficients to the remaining components, we obtained the denoised data. Finally, we re-referenced the denoised data to the average activity of all electrodes.

### Inter-subject correlation

To evaluate whether the evoked responses to the stimuli were shared among participants, we calculated the ISC of neural responses to the video ads by calculating the correlation of EEG activity among participants. Researchers have established that, for an evoked response across participants (or trials) to be reproduceable, it is necessary that each participant (trial) provides a reliable response [11]. In this sense, ISC is similar to traditional methods that capture the reliability of a response by measuring the increase in magnitude of neural activity; the important difference is that, by measuring reliability across participants, we avoid presenting stimuli multiple times to a single participant. In addition, this method is compatible with continuous naturalistic stimuli such as video ads. Therefore, the goal of ISC analysis is to identify correlated EEG components that are maximally shared across participants. By “EEG component,” we mean a linear combination of electrodes, which can be considered “virtual sources” [11]. This correlated component analysis method is similar to principal component analysis, for which, instead of the maximum variance within a dataset, the maximum correlation among datasets is considered. Like other component extraction methods, this method identifies components by solving a eigenvalue problem [40,41]. Below, we explain our ISC calculation procedure for multiple stimuli (provided in [11,22]).

To construct the input data, we combined data from participants in all four conditions (six videos for each condition; see Fig 6-a for one stimulus) and obtained 24 three-dimensional EEG matrices (channel × data samples × participants). For each of these 24 matrices, we separately calculated the between-subject cross-covariance using Eq1:

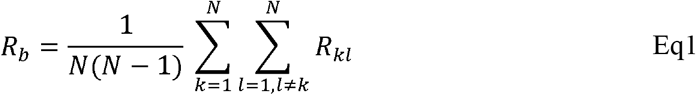

where *R_kl_* is as follows:

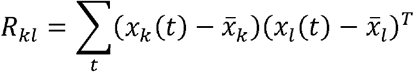

*R_kl_* indicates cross-covariance among all electrodes of subject *k* with all electrodes of subject l. The matrix *x_k_*(*t*) contains 32 electrode activities (preprocessed data) of subject *k* measured in time, and 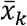 is the average of *x_k_*(*t*) over time. Additionally, we separately calculated the within-subject cross-covariance for each of the abovementioned matrices using Eq2:

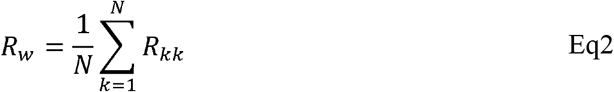

*R_kk_* is calculated in an identical manner to *R_kl_*, except it considers only the electrode activity of subject *k*. We then summed the calculated *R_b_* and *R_w_* of all 24 matrices to obtain the pooled within- and between-subject cross-covariances, representing data on all stimuli.

**Fig 6:**
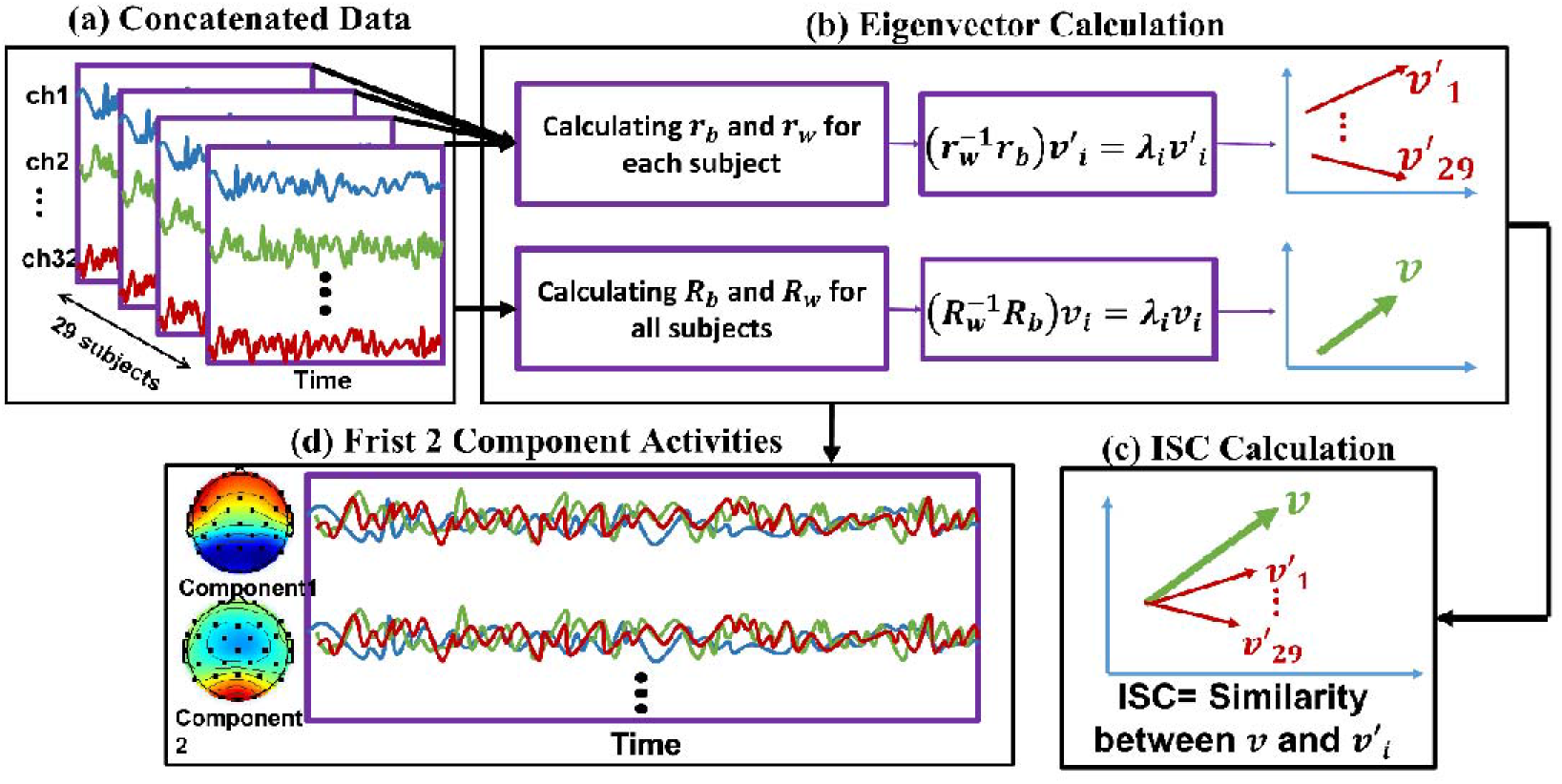
The ISC calculation and component activity estimation procedure. We repeated this procedure separately for each stimulus. **(a)** We first concatenated the EEG activity of all participants together. **(b: upper line)** For each subject, we calculated the and (using Eq1 and Eq2, respectively) based on the cross-covariance matrices. By solving the eigenvalue problem (Eq3), we computed eigenvectors of each subject. The red arrows represent each subject’s eigenvector (). Therefore, we obtained 29, corresponding to the number of participants. Note that each is a 32-dimensional matrix corresponding to the number of electrodes (for simplicity, the vectors are shown in two dimensions). **(b: lower line)** We summed all the across all participants to calculate and did the same for to calculate. Then, by solving the eigenvalue problem, we obtained, which represents maximal correlation across participants. Note that is a 32-dimensional matrix corresponding to the number of electrodes (for simplicity, the vectors are shown in two dimensions). **(c)** We calculated the similarity between the representative vector of all participants () and each of the vectors corresponding to each subject () to obtain ISC. **(d)** Using the calculated eigenvectors and forward model, we calculated the scalp activity of each component, which are linear combinations of electrode activity (32 components were calculated with the order from strongest to weakest). In sum, the activity of first two strongest components is displayed here.

Then, to improve the robustness of our analysis against outliers, we used the shrinkage regularization method to regularize the within subject-correlation matrix[42] (Eq3):

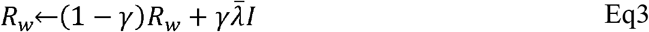

where 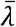 is the average of eigenvectors of *R_w_*, with *γ* equal to 0.5.

Next, by solving the eigenvalue problem for 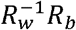 (Eq4), we obtained eigenvectors *v_i_*. Such eigenvectors are projections indicating the maximum correlation among participants, from strongest to weakest, provided by eigenvalues *λ_i_*.

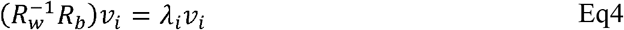

After this step, we obtained the projections that are maximally correlated among all participants considering all stimuli (Fig 6-b). Next, to measure the reliability of individual participants’ EEG responses, we calculated the correlation of projected data for each subject with projected data for the group separately for each stimulus (Fig 6-c). This metric shows how similar the brain activity of a single subject was to that of all other participants. Next, we calculated the ISC using the correlation of such projections for each stimulus (video ads) and component, averaged across all possible combinations of participants. Thus, for each stimulus and each subject, we obtained a component matrix. By summing the first two strongest components, we obtained the ISC for each subject, as follows (Eq5):

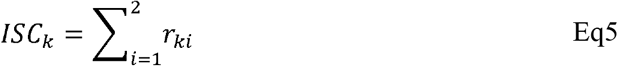

where

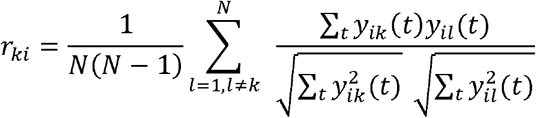

*r_ki_* is the Pearson correlation coefficient averaged across all pairs of participants applied to component projection *y_ik_*, which is defined as follows (Eq6):

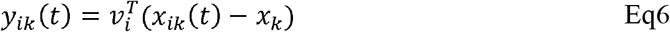

Next, we summed the first two strongest components to represent the ISC of each subject for each stimulus (*i* =1:2) and ignored the weaker components because they were below chance level in one condition. Finally, we averaged the calculated ISC over the six stimuli of each condition to obtain the ISC of all participants in each of the four conditions. To illustrate the scalp activity of each component, we used the corresponding forward model following previous studies [43,44] (Fig 6-d). In Fig 2-a, the bars illustrate the sum of the two first components.

To determine chance-level ISC, we first built up phase-randomized EEG data using a method that randomizes EEG signal phases in the frequency domain [11,45]. Using this method, we obtained new time series where the temporal alterations were not necessarily aligned with the original signals and therefore not correlated across participants. We then implemented the aforementioned steps on the randomized data, identically: for each condition, we computed within- and between-subject cross-covariances, projected the data on eigenvectors, calculated ISC, and averaged them over stimuli. We generated 100 sets of such randomized data and continued the process as described above. Finally, we obtained the chance-level ISC by averaging the values computed in these 100 iterations. The results attributed to chance level are shown by the gray bars in Fig 2-b.

### ISC-HL

We also tested whether we could assign a single participant to the HL or LL group based on participant’s neural brain activity. In other words, we tested whether we could predict the level of narrativity the participant was exposed based on EEG activity when participants attended to HL and LL videos. To do so, we calculated the projections v_i_ using only the data from the HL group. We then calculated the ISC of each participant (from both the HL and LL groups) based on the projections calculated only from the HL group and called the result “ISC-HL.” Therefore, in each group, each participant had an ISC-HL value showing how similar their neural activity was to the activity of all participants in the HL group. The difference between ISC-HL and ISC is that, while calculating ISC, we explored how similar the activity of a participant was to their own group (whether HL or LL). When calculating ISC-HL, however, we calculated how similar the activity of participants in both groups (HL and LL) was to the activity of only the participants in the HL group. We hypothesized that the neural activity of a single subject from the HL group would be more similar to the overall activity of all participants in HL group (i.e., higher ISC-HL) than to the neural activity of a single subject from the LL group. To avoid bias in this procedure, we excluded the test subject from the HL group while computing the projections *v_i_* and then calculated the ISC-HL for the corresponding subject. As described in the previous section, we summed the two first stronger components of ISC-HL and also averaged them across six stimuli for each condition. Additionally, we assessed classification performance for both groups (HL vs. LL) using the AUC characteristics of a fitted support vector machine model with 10-fold cross-validation on the data. To determine the chance-level AUC, we randomly shuffled the labels 1,000 times. In each iteration, AUC was calculated through an identical process, starting from extracting the correlated components of HL-labeled group and following all subsequent steps described above. We then determined the significance level by comparing the actual AUC value computed from the correct labels to the average of 1,000 randomized AUC values. We repeated all the aforementioned steps separately for the PC and VR conditions.

### Statistical analysis

To test statistical differences across narrativity levels, we conducted a two-way ANOVA. Based on the study design, our independent variables were narrativity level (HL vs. LL) and medium (PC vs. VR). The dependent variables were the scores from the two questionnaires (i.e., attention and immersion) and calculated neural reliability (as expressed by ISC). The output of such a comparison includes the main effects of “narrativity level” and “medium” and the interaction effect of “narrativity level × medium.” The same procedure and dependent variables, but with narrativity level (HL vs. LL) and viewing order (first vs. second) as independent variables, were used to test statistical differences across viewing order. In the present study, we focused on the effect of narrativity level and viewing order on brain activity patterns by comparing the calculated features in HL vs. LL conditions, and first vs. second viewing. Therefore, we do not report results for the main effect of medium. Nevertheless, we included medium type as a factor in the statistical analyses to control for its effect. Additionally, we performed post-hoc analysis when we identified interaction effects but also when we judged it appropriate to report simple effects. We corrected the p-values of the post-hoc tests using the Bonferroni method. Finally, to avoid multiple comparison error of multiple statistical tests (such as comparing HL and LL within two extracted components; see results), we corrected the p-values based on the number of statistical test repetition errors (FDR correction) using the Benjamini-Hochberg method [46]. To test the role of engagement in the relationship between narrativity level and ISC, we performed a mediation analysis: narrativity level → engagement → ISC. The analysis was performed using model 1 of the MEMORE V2.1.2 macro in SPSS version V28 [47]. This model represents a single variable (i.e., the mediator) that is proposed to mediate a relationship between one independent variable and the dependent variable. The analysis estimates the impacts of the total, direct, and indirect effects of the independent variable (i.e., narrativity level) on the dependent variable (i.e., ISC). The total effect represents the effect of the narrativity level on ISC without considering the mediator (i.e., engagement). The direct effect represents the effect of narrativity level on ISC controlling for the effect of engagement. The indirect effect represents the mediation effect of narrativity level on ISC via engagement. The 95% confidence intervals were calculated using a percentile bootstrap with 5,000 samples.

## Acknowledgements

We are grateful to Tirdad Seifi Ala for helping with technical aspects of the data analysis and for revising an early version of the manuscript, and to Thomas Anthony Pedersen for providing technical support and help during the early stages of the study.

